# The Multiomics Blueprint of Extreme Human Lifespan

**DOI:** 10.1101/2025.02.24.639740

**Authors:** Eloy Santos-Pujol, Aleix Noguera-Castells, Marta Casado-Pelaez, Carlos A. García-Prieto, Claudia Vasallo, Ignacio Campillo-Marcos, Carlos Quero-Dotor, Eva Crespo-García, Alberto Bueno-Costa, Fernando Setién, Gerardo Ferrer, Veronica Davalos, Elisabeta Mereu, Raquel Pluvinet, Carles Arribas, Carolina de la Torre, Francisco Villavicencio, Lauro Sumoy, Isabel Granada, Natalie S. Coles, Pamela Acha, Francesc Solé, Mar Mallo, Caterina Mata, Sara Peregrina, Toni Gabaldón, Marc Llirós, Meritxell Pujolassos, Robert Carreras-Torres, Aleix Lluansí, Librado Jesús García-Gil, Xavier Aldeguer, Sara Samino, Pol Torné, Josep Ribalta, Montse Guardiola, Núria Amigó, Oscar Yanes, Paula Martínez, Raúl Sánchez-Vázquez, Maria A. Blasco, Jose Oviedo, Bernardo Lemos, Julia Rius-Bonet, Marta Torrubiano, Marta Massip-Salcedo, Kamal A. Khidir, Thong Huy Cao, Paulene A. Quinn, Donald J. L. Jones, Salvador Macip, Eva Brigos-Barril, Mauricio Moldes, Fabio Bartieri, Gerard Muntané, Hafid Laayouni, Arcadi Navarro, Manel Esteller

## Abstract

The indexed individual, from now on termed M116, was the world’s oldest verified living person from January 17^th^ 2023 until her passing on August 19^th^ 2024, reaching the age of 117 years and 168 days (https://www.supercentenarian.com/records.html). She was a Caucasian woman born on March 4^th^ 1907 in San Francisco, USA, from Spanish parents and settled in Spain since she was 8. A timeline of her life events and her genealogical tree are shown in **Supplementary Fig. 1a-b**. Although centenarians are becoming more common in the demographics of human populations, the so-called supercentenarians (over 110 years old) are still a rarity. In Catalonia, the historic nation where M116 lived, the life-expectancy for women is 86 years, so she exceeded the average by more than 30 years (https://www.idescat.cat). In a similar manner to premature aging syndromes, such as Hutchinson-Gilford Progeria and Werner syndrome, which can provide relevant clues about the mechanisms of aging, the study of supercentenarians might also shed light on the pathways involved in lifespan. To unfold the biological properties exhibited by such a remarkable human being, we developed a comprehensive multiomics analysis of her genomic, transcriptomic, metabolomic, proteomic, microbiomic and epigenomic landscapes in different tissues, as depicted in **Fig. 1a**, comparing the results with those observed in non-supercentenarian populations. The picture that emerges from our study shows that extremely advanced age and poor health are not intrinsically linked and that both processes can be distinguished and dissected at the molecular level.

**Fig. 1.**
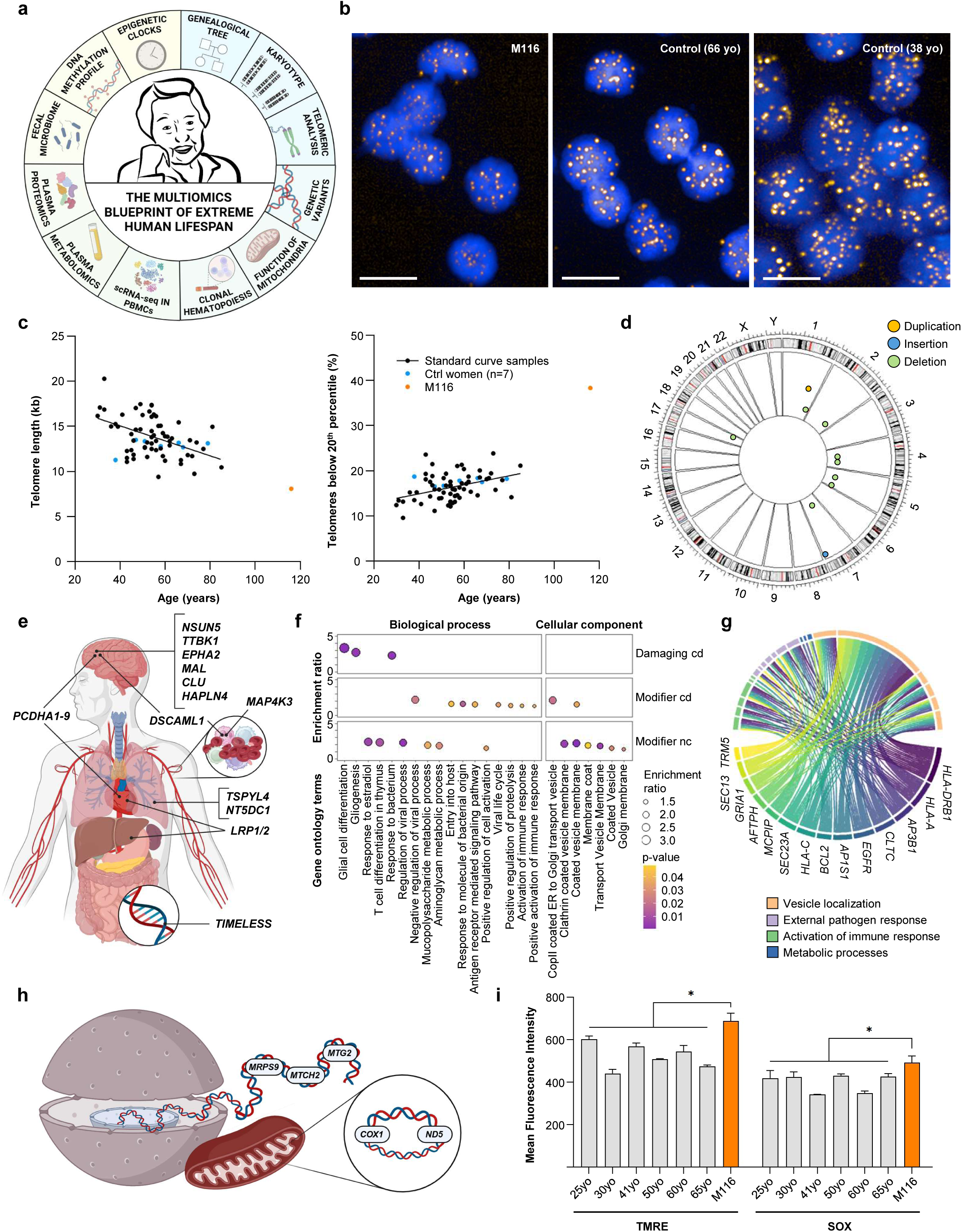
Chromosomes and genes. **a**, Schematic representation of all –omics studied in the supercentenarian. **b**, Telomeres marked with Cy3 (yellow) in nuclei stained with DAPI (blue) observed in HT-qFISH from M116 and younger women’s PBMCs. Scale bars: 20 µm. **c**, Telomere length (Kb) calculation (left) and percentage of extremely short telomeres (below the 20th percentile) (right) in M116 (orange) using standard curve from samples previously analyzed (black) and control women (blue) (**Online Methods**). **d**, Circos plot with chromosomal alterations detected through optical genome mapping in supercentenarian. **e**, Variants of interest (VOI)-harboring genes found in supercentenarian’s genomic DNA contributing to immune function, cardiovascular health, neuroprotection, metabolism, and DNA dynamics. **f**, Significantly enriched functions of VOI-harboring genes in the supercentenarian. **g**, VOI-harbouring genes significantly contributing to enriched functions. **h**, VOI-harboring genes found in supercentenarian’s genomic and mitochondrial DNA contributing to mitochondrial function. **i**, Mean fluorescence intensity of TMRE (a marker of mitochondrial membrane potential) and SOX (a marker of mitochondrial superoxide ion) in PBMCs from the supercentenarian (orange) and healthy controls across various ages (gray). Unpaired t-test was used to statistically compare M116 to the mean of all control women. *p < 0.05.

## RESULTS AND DISCUSSION

Samples from the subject were obtained from four different sources: total peripheral blood, saliva, urine and stool at different times. Most of the analyses were performed in the blood material at the time point of 116 years and 74 days, unless otherwise specifically indicated (**Data set 1**). The simple karyotype of the supercentenarian did not show any gross chromosomal alteration (**Supplementary Fig. 1c**). Since many reports indicate the involvement of telomeres in aging and lifespan^1^, we interrogated the telomere length of the M116 individual using High-Throughput Quantitative Fluorescence In Situ Hybridization (HT-Q-FISH) analysis^2^. Illustrative confocal images with DAPI staining and the telomeric probe (TTAGGG) for M116 and two control samples are shown in **Fig. 1b**. Strikingly, we observed that the supercentenarian exhibited the shortest mean telomere length among all healthy volunteers^3^ with a value of barely 8 kb (**Fig. 1c**). Even more noticeably, the M116 individual displayed a 40% of short telomeres below the 20^th^ percentile of all the studied samples (**Fig. 1c**). Thus, the observed far reach longevity of our case occurred in the chromosomal context of extremely short telomeres. Interestingly, because the M116 individual presented an overall good health status, it is tempting to speculate that, in this setting, telomere attrition behaves more as a chromosomal clock for aging rather than a predictor of age-linked diseases such as neurodegeneration or diabetes. As food for thought, the huge erosion of the telomere sequences in the supercentenarian could also relate to the absence of any diagnosed cancer because it could limit the replicate lifespan of any malignant cell.

Beyond telomeric regions, we assessed the presence of structural variants (SVs) at the chromosomal scale that could not be detected by regular karyotyping. Using Optical Genome Mapping (OGM) (**Methods Online**), we observed the presence of ten rare SVs in M116 as described in **Data set 2.** The largest SVs were a 3312.4 kb deletion in chromosome 4 and a 93.5 kb deletion in chromosome 17, as illustrated in the Circos plot of **Fig. 1d**. Extreme longevity is believed to be influenced by a complex interplay of genetic variants, both common and rare, that affect healthy aging. This study sought to investigate rare genetic variants in M116’s genome that may contribute to her exceptional lifespan. We performed Whole Genome Sequencing (WGS) of the M116 individual where we identified approximately 3.8 million single nucleotide variants (SNVs) across her genome. Variants were filtered and annotated, and rare variants were defined as having a variant allele frequency (VAF) below 0.015 in European populations (1000 Genomes and gnomAD) (**Online Methods**). Then, they were further annotated attending to their potential impact on protein function, and if applicable, classified as variants of interest (VOI) (**Online Methods**), ending up with 91,666 VOI affecting 25,146 genes. The analysis focused on identifying functional variants that might impact genes or gene sets associated with longevity or disease resistance. The comparison was made between M116’s genome and a control set of 75 Iberian women from the 1000 Genomes Project to identify “extreme” variants potentially linked to her longevity. We identified 7 homozygous variants in M116’s genome, affecting 16 protein-coding and 3 non-coding genes (**Data set 3**). Remarkably, none of these rare homozygous variants were found in the control European populations, suggesting that these variants could be unique to her and may contribute to her exceptional longevity. Examples include homozygous variants detected on *DSCAML1*, a gene associated with immune function and cognition retention^4^; *MAP4K3*, linked to lifespan regulation of *Caenorhabditis elegans*^5^ and to autoimmune disease, cancer and aging^6^; *TSPYL4* and *NT5DC1*, linked to homeostatic pulmonary function^7^ and the protocadherin alpha cluster (*PCDHA1-9*), related to aging brain health and heart disease^8,9^ (**Fig. 1e and Data set 3**).

We also conducted an overrepresentation of functional categories analysis to evaluate whether specific biological processes or pathways were disproportionately affected by the rare variants in M116’s genome. In addition to glial functionality—potentially linked to neuroprotection against degeneration—and the classic hormonal factors associated with aging associated features (“Response to Estradiol”)^10^, one of the strongest enrichments was found in immune system-related categories, such as “T Cell Differentiation in thymus”, “Response to Bacterium”, “Regulation of Viral Process” or “Antigen Receptor Mediated Signalling Pathway” (**Fig. 1f** and **Data set 4**). Furthermore, we observed VOI affected genes with highly pleiotropic functions across several overrepresented immune system-related pathways (**Fig. 1g**). These functions are essential for controlling infections, autoimmune regulation, and possibly cancer surveillance, which could have contributed to M116’s longevity. Categories related to heart function, such as “Atrial Septum Morphogenesis” and “Myocardial Fibrosis“; cholesterol metabolism and insulin signalling; and neuroprotection and axonal function were also significantly enriched (**Data set 4**). In this regard, rare variants were found in genes involved in lipid metabolism and heart function, such as *LRP1* and *LRP2*^11^, and in *NSUN5* and *TTBK1*, genes linked to neuroprotection that could potentially be contributing to the preservation of cognitive function in extreme old age (**Fig. 1e**). Additionally, we found a rare variant in the *TIMELESS* DNA repair gene, linked to the evolutionary divergence in longevity across *Drosophila* populations^12^ (**Fig. 1e**). Interestingly, due to the long studied effect of reactive oxygen species in aging, rare variants in many genes involved in mitochondrial function, such as *ND5*, *COX1*, *MTG2*, *MTCH2*, and *MRPS9*, were detected (**Fig. 1h**). These variants could affect mitochondrial oxidative phosphorylation, a process crucial for energy production, aging and longevity^13^. To experimentally interrogate this possibility, peripheral blood mononuclear cells (PBMCs) from M116 were isolated to assess mitochondrial membrane potential and to measure superoxide ion levels (**Online Methods**). We observed that the values of the markers were even higher than the ones observed in PBMCs from younger women (**Fig. 1i**), indicating not only preserved but robust mitochondrial function in the supercentenarian. Finally, we conducted three burden tests to identify genes or gene sets with a significantly higher burden of rare variants in M116 compared to controls, which were considered as differentiating genes, and performed an over-representation analysis (**Online Methods**). Using this approach, key rare variants in genes involved in neuroprotection and longevity, such as *EPHA2, MAL*, *CLU* and *HAPLN4* were also highlighted (**Fig. 1e and Data set 5**). Overall, our study indicates that there was not a single biological process targeted by a unique rare genetic variant solely associated with the healthy aging and extended lifespan of M116, but rather a combination of rare variants in multiple genes and pathways (immune system, cardioprotection, brain activity and mitochondrial metabolism) likely working together that promoted her remarkable longevity.

To dwell further in the molecular and cellular landscapes associated with the extreme lifespan of the individual M116, we studied in deep detail the characteristics of the subject’s blood. This is an important issue since this source of biological material is the one commonly used in the laboratories of the hospitals around the world; and there are exciting data from parabiotic experiments in mice suggesting that sharing bloodstream can affect aging^14^. We first assessed the existence of clonal hematopoiesis of indeterminate potential (CHIP), also denominated age-related clonal hematopoiesis that it is associated with the advanced age^15^. The occurrence of CHIP is thought to be a precursor of myeloid malignancies and other cancers and fosters the development of cardiovascular disease^15^. Defining CHIP by the presence of mutations in any leukemia driver gene (**Data set 6**) with a VAF ≥2%, we found that M116 harbored a CHIP genotype defined by one mutation in the RNA splicing factor *SF3B1* and two different mutations in the DNA demethylase *TET2* (**Fig. 2a**). Importantly, despite these predisposing mutations, the supercentenarian did not experience any tumorigenic process or cardiovascular disorder in her lifetime. We next characterized the distinct immune populations in M116 by analyzing her PBMCs using single-cell transcriptomics (scRNA-seq). This approach enabled the identification of major lymphoid and myeloid populations, including naive and memory B cells, NK cells, classical and non-classical monocytes, and distinct T cell subpopulations. The distribution of these populations is visualized in the UMAP plot in **Fig. 2b**, where each cell cluster is colored according to its cell type annotation. A striking feature is the presence of an expanded cluster of age-associated B cells (ABCs). This subpopulation is known to accumulate with age, as well as in several immunological disorders. The ABC cluster in the M116 individual is distinguished by the high expression of ABC associated markers, such as *SOX5* and *FCRL2*. We next sought to compare the immune cell type proportions of the M116 case with those of healthy individuals spanning different age groups, from young adults (25 years) to other supercentenarians. **Fig. 2c** illustrates the relative abundance of immune cell populations across five previously characterized age groups^16^, seven supercentenarians^17^, and the oldest supercentenarian M116. In younger age groups, the immune profile is characterized by higher fractions of regulatory and naive T cells. In supercentenarians, these populations are reduced and accompanied by an expansion of other lymphoid populations. Notably, in our longest-living supercentenarian M116, effector and memory T cells dominate the T cell compartment, particularly cytotoxic T cell subsets. This pattern aligns with previous observations of cytotoxic T cell enrichment in supercentenarians^17^. However, the pronounced expansion of the ABC cluster in the M116 individual is not observed in other supercentenarians, illustrating the singularity of her exceptional longevity. The observed occurrence of CHIP and the unique scRNA-seq landscape of the blood in M116 prompted us to study further the molecular composition of this clinically applicable biological source. To achieve this aim, we performed whole proton nuclear magnetic resonance (^1^H-NMR) analyses to obtain the lipid and lipoprotein landscape, the glycoprotein profile and the Low Molecular Weight Metabolites (LMWM) spectrum (**Online Methods**). All the values derived from these analyses are shown in **Supplementary Fig. 2**. The lipoprotein and glycoprotein values derived for the supercentenarian were compared with 6,022 individuals of two Spanish population-base cohorts^18,19^, thus, with similar background to M116. For the comparison of concentrations in aqueous metabolome and lipid families we used the 1,965 individuals of the Di@betes Study^18^, where these data are available. The unfolded metabolomic landscape of the oldest human reflects an interesting dichotomy: she showed values of many metabolites and biomolecules associated with a healthy life and having had a long lifespan, but at the same time a few biomarkers were suggesting that the end of her life was near. Probably the most relevant finding was that she presented one of the most efficient lipid metabolisms reported, a trait that the UK Biobank publications link to extended longevity^20^ and absence of dementia^21^, as it occurred in the M116 subject. In this regard, M116 displayed extremely low levels of VLDL-cholesterol and triglycerides, whereas HDL-cholesterol (the *“good”* cholesterol) was very high (**Fig. 2d**). Additionally, the high number of medium and large HDL-cholesterol particles and large LDL-cholesterol particles, combined with a low number of small HDL-cholesterol particles, support the idea of effective lipoprotein maturation and overall enhanced lipid metabolism^22^ (**Fig. 2d**). The supercentenarian’s highly efficient lipid metabolism is also reflected in the low levels of other lipid-related biomarkers typically associated with poor health and increased mortality, such as saturated fatty acids^23^, esterified cholesterol^23^, linoleic acid^22^, and acetone^20^, as well as in the high levels of free cholesterol^22^, associated with good health and survival (**Fig. 2d**). Beyond the lipid metabolism, the M116 supercentenarian exhibited low concentrations of glycoproteins A and B (**Fig. 2d**) indicating the existence of a healthy inflammatory profile that precludes the occurrence of evident systemic inflammatory diseases and associated inflammaging phenotypes with clinical impact^24^. In particular, metabolites associated with cardiovascular risk support the idea of an excellent cardiovascular health in the supercentenarian **(Fig. 2e)**. Interestingly, the only metabolites that suggest her real very advanced chronological age and the risk of death are the low levels of the aminoacids glycine, histidine, valine, and leucine, and the high concentration of lactate and creatinine^20^ (**Fig. 2d**). Overall, these metabolomic findings suggest that a highly engaged lipid metabolism together with very low levels of inflammation could explain the excellent health and extreme longevity observed in the studied supercentenarian, despite signs of functional decay in other pathways.

**Fig. 2.**
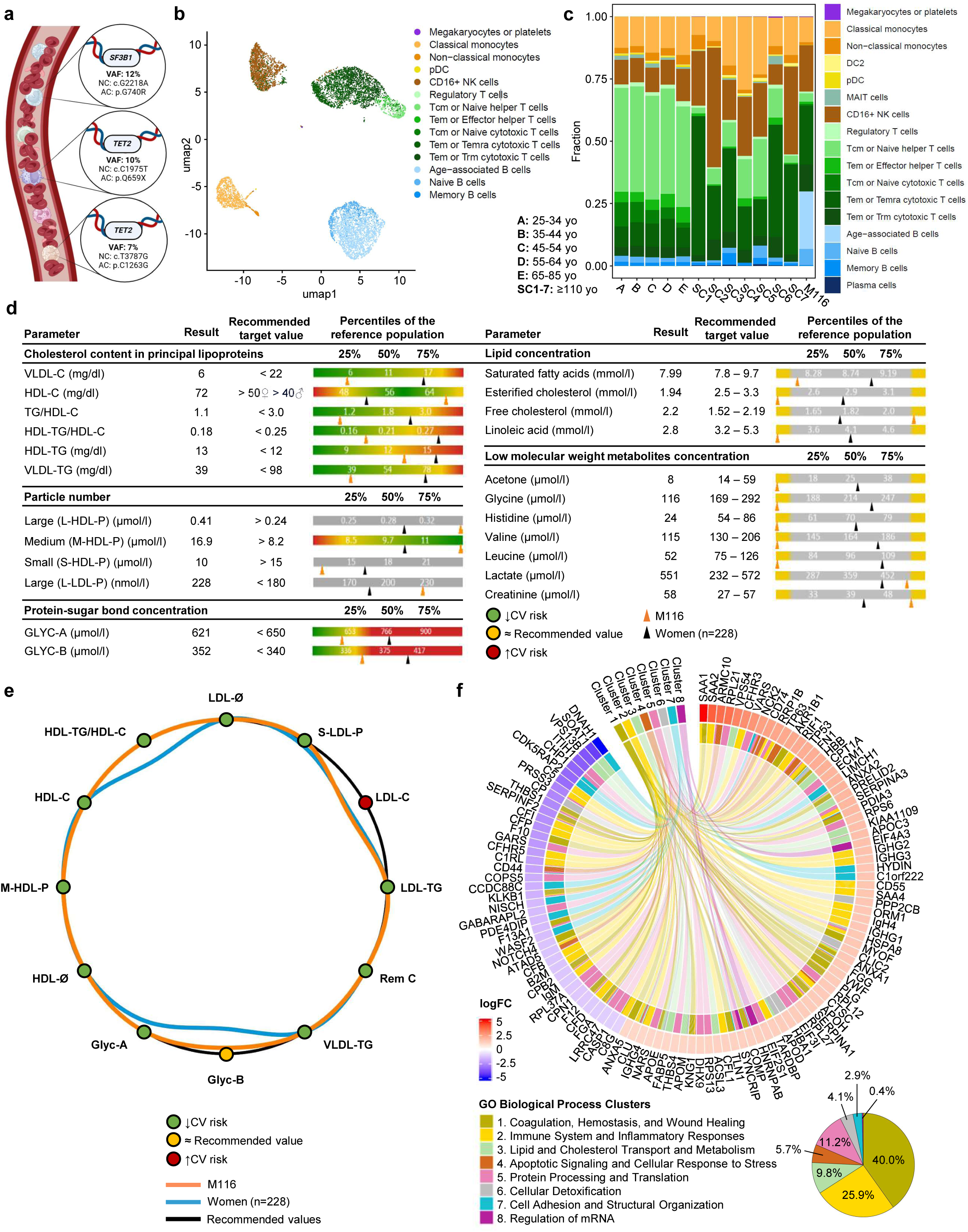
Blood genomics, metabolomics and proteomics. **a**, Mutated genes contributing to supercentenarian’s clonal hematopoiesis of indeterminate potential (CHIP). VAF: variant allele frequency; NC: nucleotide change in cDNA; AC: aminoacid change in protein. **b**, UMAP of PBMCs from supercentenarian coloured by cell type annotation after single-cell RNA-seq. **c**, PBMCs’ cell type proportions comparison of M116 with healthy controls of five different age ranges and other previously studied supercentenarians (**Online Methods**). **d**, Characteristic metabolic signatures in supercentenarian’s plasma represented alongside recommended target values established from reference population. Percentiles in reference population are represented in bars, where arrows indicate levels in supercentenarian (orange) and control population (black). Those variables clearly associated with cardiovascular (CV) risk appear in a colour scale, with green representing low CV risk and red representing high CV risk. **e**, Lipidic contour of supercentenarian (orange) and control population (blue) modelling 12 lipid and inflammatory metabolism variables associated with cardiovascular (CV) risk. Recommended values in black (**Online Methods**). **f**, Proteomics data showing differentially expressed proteins between M116 and controls and their associated functional category (p-value and q-value < 0.05, with at least three associated proteins per functional category). Protein colors range from red (upregulated) to blue (downregulated), according to fold change in expression. 1-8 gene ontology (GO) biological process clusters (colored) were obtained from clustering GO terms based on semantic similarity (**Online Methods**).

Since the composition of extracellular vesicles (ECV) has also been linked to longevity^25^, we performed an additional proteomic analysis of M116 ECVs and compared it to a baseline of younger post-menopausal women (yPM; n=4, ages: 49-65 years). Using an Orthogonal Partial Least Squares Discriminant Analysis (OPLS-DA) model (**Online Methods**), we observed proteomic distinctions between both groups (**Supplementary Fig. 3**). Overall, 231 proteins had significant expression differences among M116 and the post-menopausal women set (**Data set 7**). Gene Ontology (GO) enrichment analysis clustered in eight groups (coagulation, immune system, lipid metabolism, apoptosis, protein processing, cellular detoxification, cellular adhesion and mRNA regulation) (**Fig. 2f**), showing an enrichment of pathways related to complement activation, B cell immunity, acute inflammatory response, adaptive immune response, humoral response and, potentially, a reduced inflammatory response in the supercentenarian (**Data set 8**). Furthermore, we also found an increased lipid and cholesterol transport, higher lipoprotein remodeling, enhanced fatty acid transport, lipoprotein clearance and high regulation of lipoprotein levels in the M116 sample (**Data set 8**); in addition to enhanced oxidant detoxification and response to oxidative stress (**Fig. 2f and Data set 8**). These differences suggest an increase in protective mechanisms that could contribute to a healthy aging in the interrogated individual. Interestingly, these protein biomarkers of well-being in the M116 individual coexisted with a potentially dangerous pathophysiological sign: the protein most elevated in the supercentenarian compared to the yPM women was serum amyloid A-1 protein (SAA1) (**Data set 7**). SAA1 is linked to Alzheimer’s disease^26^, but despite her advanced age, M116 showed no evidence of any neurodegenerative disorder.

The above discoveries drove us to learn about the microbiome composition of the studied record-breaking supercentenarian since microorganisms are critical in determining not only the metabolite composition of our body, but also inflammation, intestinal permeability, cognition and bone and muscle health^27^. Thus, we determined the fecal microbiota composition by 16S rDNA analysis of the M116 individual and compared the results with 445 samples from control individuals (250 women and 195 men) aged 61-91 years that were not under antibiotic treatment from the curatedMetagenomicData data set (**Online Methods**). First, we analyzed the microbiome diversity within the samples from M116 and compared it to that of control women individuals. Focusing on the α-diversity, a measure of diversity within-sample, we observed that the M116 sample exhibited a higher value compared to the mean alpha-diversity of the control female population (Shannon alpha-diversity values: 6.78 vs. 3.05, respectively). Next, we examined the beta-diversity, which measures dissimilarity between conditions. As shown in the PCA plot (**Supplementary Fig. 4**), the M116 sample is positioned far from the cluster of control samples, emphasizing the distinctiveness of her microbiome compared to the controls. At the phylum level, the most startling finding was the high levels of *Actinobacteriota* in comparison with the control populations from both genders (**Fig. 3a**). Zooming at the family level we observed that the increase was mostly due to the elevated amount of *Bifidobacteriaceae* (**Fig. 3b**), particularly *Bifidobacterium* when we move to the genus classification (**Fig. 3c** and **Fig. 3d**). This finding contrasts sharply with the typical decline of this bacterial genus in older individuals^28^; however, it has also been reported at elevated levels in centenarians^29^. *Bifidobacterium* is thought to be a beneficial bacterium contributing, among other processes, to anti-inflammatory responses, an observation that links with the low levels of inflammation markers in the metabolomics study (**Fig. 2d**). High content of *Bifidobacterium* has also been associated with the production of short-chain fatty acids and conjugated linoleic acid^30^, observations that relate to the “healthy” lipid-related biomarker profile detected by ^1^H-NMR (**Fig. 2d** and **Fig. 2e**). Importantly, the use of *Bifidobacterium* as a probiotic that could slow down the progression of many aging-associated disorders is gaining momentum. Intriguingly, our healthy supercentenarian ingested around 3 yogurts every day containing *Streptococcus thermophilus* and *Lactobacillus delbrueckii subsp. bulgaricus*, known to favor the growth of the described bacteria in the gut (a representative 3-weeks menu of the diet that the supercentenarian followed during her last 20 years of life is shown in **Data set 9**). Thus, this could be an example of a dietary intervention that, acting in the gut microbiota, is associated with healthy aging and long lifespan. Other bacteria might also contribute to the described phenotype. For example, lower levels of the phylum *Proteobacteria* and *Verrucomicrobiota,* as we have found in our M116 sample vs the control groups (**Fig. 3a**), have been linked to older adults without age-related frailty^31^. In a similar manner, the diminished levels of the pro-inflammatory genus *Clostridium*^28^ in the supercentenarian vs controls (**Fig. 3d**) would support the absence of aging-associated inflammatory diseases. Overall, these results suggest that the studied extreme supercentenarian possesses a microbiome that confers an increased likelihood for a healthy extended lifespan. This is also consistent with her adherence to a Mediterranean diet (**Data set 9**), which might have contributed to the described unique microbiome composition^32^.

**Fig. 3.**
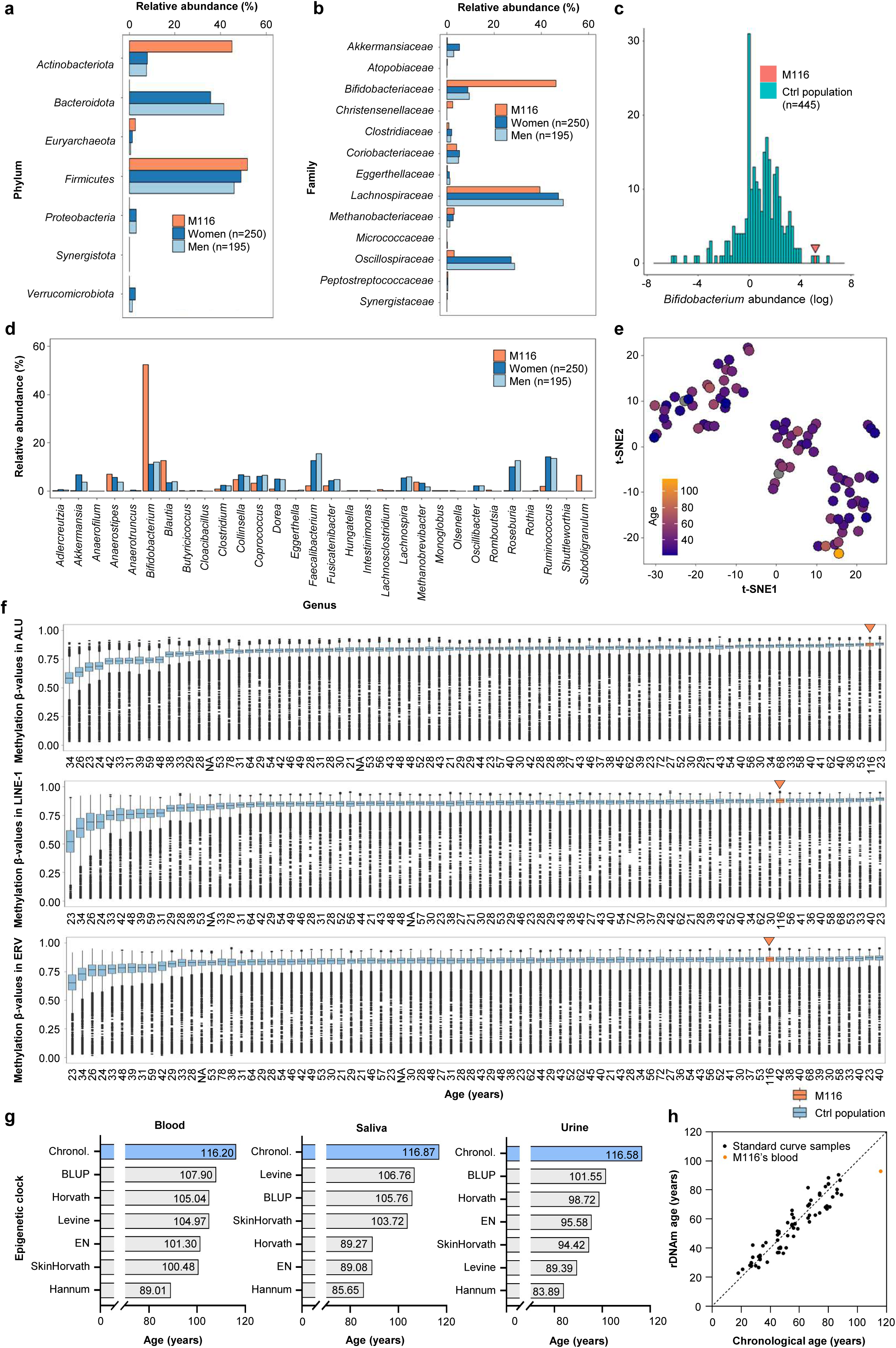
Microbiome and epigenetics. **a-c**, Bar plots representing the percentage of relative abundance of bacterial populations found in supercentenarian’s faecal samples (**a**, at Phylum level; **b**, at Family level; **c**, at Genus level). **d**, Abundance of *Bifidobacterium* in supercentenarian (orange) and the control population (blue) represented as the count of individuals. Orange arrow points to supercentenarian. **e**, tSNE representation of differentially methylated CpGs between supercentenarian and healthy control population (**Online Methods**). **f**, Boxplot representing predicted methylation values in ALU, LINE-1, and ERV repetitive element sequences for supercentenarian (orange) and healthy control population (blue). Orange arrows point to supercentenarian. Boxplots are ordered increasingly according to median methylation value for each sample. **g**, Chronological (blue) and estimated biological age (gray) of supercentenarian’s tissues according to different publicly available epigenetic clocks. **h**, Estimated rDNA methylation age in supercentenarian’s blood (orange) and control population (black) according to rDNAm clock (**Online Methods**).

Finally, we added the layer of the epigenetic setting in the studied longest-living human, as a factor (like microbiome composition) that might provide a dynamic, plastic and adaptable read-out of M116 biology. DNA methylation is probably the most studied epigenetic mark in cell biology and disease^33^, being also disrupted as we age^34^, as it has been shown, for example, comparing newborns vs nonagenarians^35^ and in Hutchinson-Gilford progeria^36^. Thus, we first interrogated the CpG methylation status of the supercentenarian case using a comprehensive microarray that analyzed more than 850K loci^37^, comparing the results to a large collection of control individuals previously studied in our laboratory^38^ (n=81, 21-78 years old) and interrogated with the same epigenetic platform (**Online Methods**). We observed that 69 CpG sites were differentially methylated (β-value at least 50% distinct) in the supercentenarian vs the rest of individuals (**Data set 10**). According to CpG content, only 26.09% (18 out of 69) of these sites were placed in CpG islands, whereas 73.91% (51 out of 69) were located in regions of the genome with lower CpG density (**Data set 11**). Related to genomic structure, 33.33% (23 out of 69) of CpG sites were located within the 5’-regulatory gene regions (Transcription Start Site -1500 base pairs, Transcription Start Site -200 base pairs, 5’-Untranslated Region and 1st exon), while the remaining 66.67% (46 out of 69) of CpG sites were associated with the gene body, 3’-Untranslated Region, or intergenic regions (**Data set 11**). Importantly, 68.12% (47 out of 69) of the differently methylated loci corresponded to a loss of CpG methylation in the supercentenarian sample. We also compared the localization of hypomethylated and hypermethylated CpG sites in relation to CpG islands and observed no remarkable differences (**Data set 11**). However, when focusing on their localization within gene loci, we found a higher proportion of hypomethylated CpG sites (21 out of 47; 44.68%) within gene bodies compared to hypermethylated CpG sites (6 out of 22; 27.27%). This loss of CpG methylation in the supercentenarian sample is in line with the previously described waves of DNA hypomethylation shifts taking place in human aging^35,36^. Interestingly, a global DNA hypomethylation status is also commonly observed in human cancer^39^, a disease with an increased incidence in older people, highlighting some of the crosstalks and commonalities between the aging and tumorigenesis pathways^1^. To provide further functional impact to the identified CpG sites, we crossed their methylation status with RNA expression data in human cell lines^40^ (**Online Methods**). For those CpG sites included in the described databases within an associated gene with available RNA values, we found that the M116 CpG hypomethylation status was associated with low expression of *EGFL7* (EGF like domain multiple 7) and *ADCY3* (adenylate cyclase 3), and also hypermethylation of *PLEKHA1* (pleckstrin homology domain containing A1) (for all cases p-value < 0.05 and rho > |0.3|) (**Data set 12**). For an additional gene, *VASN* (vasorin), the hypomethylation event at the supercentenarian occurred in the 5’-UTR and, as expected for this regulatory region^41^, was associated with high levels of the RNA transcript (**Data set 12**). The identified DNA methylation-controlled genes are *bona fide* candidates to play a role in the supercentenarian’s biology since they are involved in vascular stemness (*EGFL7*), body mass index (*ADCY3*), macular degeneration (*PLEKHA1*) and bone turnover (*VASN*). All of these activities considered hallmarks of aging or tightened to aging-associated disorders. Using the 69 differentially methylated loci in a supervised hierarchical clustering, we can observe the distinct DNA methylation profile between M116 and the controls (**Supplementary Fig. 5**). As described above, most of these differential CpG sites represented hypomethylation events in M116 compared with the other samples (**Supplementary Fig. 5** and **Data set 11**). Further dimensionality reduction analysis with the 69 differentially methylated CpG sites by t-Distributed Stochastic Neighbor Embedding (t-SNE) also shows the supercentenarian as a DNA methylome outlier (**Fig. 3e**). Importantly, because the loss of DNA methylation in repetitive sequences has been reported through the aging process^42^, we carefully studied the DNA methylation content of three families of repeats: LINE-1, ALU and ERV. Interestingly, we found that the supercentenarian did not undergo major hypomethylation events in these loci, instead retained hypermethylated CpGs even at higher levels than most of the younger individuals (**Fig. 3f**). Thus, these findings suggest that a disruption of the DNA methylation balance (hypermethylation/hypomethylation) in gene 5’-regulatory CpG islands is linked to the aging trajectory, but keeping epigenetically silent DNA repetitive sequences could confer an advantage associated with healthy longevity, as it occurred in our case.

DNA methylation analyses provided another important clue that might explain the amazing lifespan of our supercentenarian. The last years have witnessed the development of the so-called “epigenetic clocks”^43^ that are utilized as a proxy to calculate ‘‘biological age’’ of a given tissue or specimen that, although in most cases is expected to match the ‘‘chronological age’” (the amount of time that has elapsed from birth), it is not always the case. In this regard, several pathologies can accelerate the process and, for example, premature biological aging determined by these epigenetic clocks occurs among carriers of viral infections^44,45^. Since these technologies use a comparable DNA methylation platform used herein, we were able to calculate the biological age of our supercentenarian using six different epigenetic clocks (**Fig. 3g** and **Online Methods**). Remarkably, all the distinct algorithms of age based on DNA methylation yielded the same result. Our supercentenarian exhibited a much younger biological age than her real chronological age and this occurred in the three different tissues analyzed (**Fig. 3g**). These results were reinforced when we used a completely different epigenetic clock: CpG methylation status of ribosomal DNA^46^ (**Fig. 3h**). Importantly, this approach does not use the mentioned DNA methylation microarray data, but Whole Genome Bisulfite Sequencing (WGBS). Thus, we carried out WGBS of our supercentenarian sample and compared the obtained data with a set of 70 cases studied using the same experimental and bioinformatic pipeline (**Online Methods**). Our results showed that whilst the control group showed concordance between both types of age, the M116 case exhibited a much lower biological age than her chronological age (**Fig. 3h**), further validating the findings of the six epigenetic clocks derived from the DNA methylation microarray test (**Fig. 3g**). Overall, these data suggest that one of the reasons that our supercentenarian reached such a world record age was that her cells “felt” or “behaved” as younger cells, with a biological age of a centenarian.

Our findings suggest that extreme human longevity may be characterized by the coexistence of two distinct and potentially unrelated sets of features within the same individual. On one hand, there are characteristic biomarkers of very advanced age, such as shortened telomeres, clonal hematopoiesis-associated mutations, or an aged B-cell population. On the other hand, there are simultaneously preserved healthy (epi)genetic and functional tissue environment traits. This last scenario is evidenced by the presence of genetic variants protective against common diseases (e.g., cardiovascular disorders, diabetes, and neurodegeneration), an efficient lipid metabolism, an anti-inflammatory gut microbiome, and an epigenome associated with chromosomal stability and decelerated epigenetic aging. All these findings illustrate how aging and disease can, under certain conditions, become decoupled, challenging the common perception that they are inextricably linked.

## Online Methods

### Sample and Data Acquisition

Peripheral blood, urine, saliva and stool were collected from the supercentenarian and age at time of sampling was annotated and considered for subsequent analyses. The protocol of the study was approved by the institutional ethics review board (PI-23-19). An extensive interview was conducted addressing her clinical history and main lifestyle habits such as sleep time, diet, exercise, and social interactions. M116 was the world’s oldest living person at the time of starting the present study, and was the 8^th^ world’s longest-living person in history on verified records (https://www.supercentenarian.com/records.html). She was a Caucasian woman born on March 4^th^ 1907 in San Francisco, USA, to Spanish parents. She moved to New Orleans, USA, in August 1910, and in 1915, when she was 8 years old, her father passed away and she moved to Barcelona, Spain. From then on, she spent her life settled in Spain. From 2001 until death, she lived in a residence for the elderly in Olot, Catalonia, Spain. In spite of several emotionally painful events during her last years of life, like her son’s death, she kept a strong physical and mental health throughout life with good sleep habits, balanced Mediterranean diet, and active social life. She largely enjoyed from quality time with family and friends, playing with dogs, reading books, growing a garden, walking, and playing the piano. She suffered from Covid-19 and chronic age-related diseases like bronchiectasis, esophagus diverticulum, and osteoarthritis, with limited movement and high dependency during her last months of life. She never suffered from other prevalent age-related diseases like cancer or neurodegenerative diseases, unlike siblings. She passed away on August 19^th^ 2024, while sleeping, at the age of 117 years, 5 months, and 18 days.

### PBMCs extraction

Freshly-collected peripheral blood was centrifuged at 2,000*g* for 6 min at room temperature (RT) to remove plasma, and the cellular fraction was diluted with sterile PBS-EDTA 2 mM. Later, peripheral blood mononuclear cells (PBMCs) were isolated by Lymphoprep density gradient centrifugation (Serumwerk Bernburg; Cat. 00122) at 800*g* for 30 min at RT (acceleration and deceleration 3/9) and transferred to new tubes. PBMCs were centrifuged at 3,200*g* for 10 min at RT, and pellets were resuspended in 90% fetal bovine serum (FBS) + 10% DMSO for cryopreservation in liquid nitrogen.

### Karyotype

Conventional G-banding karyotype was carried out by standard procedures. Briefly, 15 million B-cells from M116 were plated in RPMI medium supplemented with 2% FBS. Colcemid was added to the culture followed by 20 minutes-incubation to arrest cells in metaphase. After centrifugation, a hypotonic solution (KCl, 0.075M) was added followed by 25 minutes-incubation at 37°C. Cells were centrifuged thrice and washed with Carnoy’s solution (methanol and glacial acetic acid, 3:1) to break cytoplasmic membranes. Chromosomes were fixed in the spindle and, along with the different fixer passes, membranes, cytoplasm, and different organelles were removed to obtain a pellet with the nuclei of the cells that were in interphase and metaphase. Finally, the G band pattern was achieved with Wright dye.

### High-Throughput Quantitative Fluorescence In Situ Hybridization (HT-Q-FISH) analysis

PBMCs (2 × 10^5^ cells per well) from M116 and control women (IJC-Campus ICO-GTP Biological Sample Collection) were plated in 96-well plate coated with poly-lysine (Greiner Bio-One, Inc.®; Cat. 655087) and fixed in Methanol: Acetic acid (3:1) for 15 minutes. Plates were stored at −20°C with the fixative solution. For Q-FISH hybridization, plates were dried at 37°C overnight and cells were then rehydrated with PBS and fixed with 4% formaldehyde, followed by digestion with pepsin/HCl and a second fixation with 4% formaldehyde. Samples were dehydrated with increasing concentrations of EtOH (70%, 90%, 100%) and incubated with the telomeric probe (TTAGGG) labelled with Cy3 at 85°C for 3 min followed by 1h at room temperature in a wet chamber. Samples were extensively washed with 50% formamide and 0.08% TBS-Tween 20 followed by TBST containing 1 μg/mL DAPI (4′,6-diamidino-2-phenylindole, dihydrochloride) (Life Technologies; Cat. D-1306) to stain the nuclei^2^. Next, the plate was washed with 0.08% TBS-Tween20 in a plate shaker for 5 min. Confocal images were captured using the Opera Phenix High-Content Screening System (PerkinElmer). Images were analyzed with Harmony High-Content Analysis Software (PerkinElmer).

### Optical Genome Mapping

High molecular weight DNA (>250 Kb) isolated from the supercentenarian B-cells was marked with Direct Label & Stain kit (DLS, Bionano Genomics®), which labels specific sequences of 6 nucleotides (CTTAAG) repeated throughout the genome using an enzyme (DLE-1) attached to a fluorophore. Images were obtained using the Saphyr instrument (Bionano Genomics®), with a minimum coverage of 300X. For result analysis, the Rare Variant Analysis algorithm included in the Bionano Solve 3.7 software, with the hg19 genome version, was used. Visualization was performed using Bionano Access 1.7.2. Alterations obtained were considered after applying the Bionano Genomics® confidence filters and discarding benign/polymorphic alterations.

### DNA extraction from peripheral blood

Freshly-collected peripheral blood was centrifuged at 2,000*g* for 6 min at RT to remove plasma, and cellular fraction was diluted with sterile PBS-EDTA 2 mM and centrifuged at 3,200*g* for 10 min at RT. Supernatant was discarded, and cells were washed twice in Ammonium-Chloride-Potassium (ACK) lysing buffer (centrifugations at 3,200*g* for 10 min at RT). Pellets with washed whole-blood leucocytes were stored at -80°C.

Lysis buffer was freshly prepared (TrisHCl 2.5 mM pH 8 + EDTA 1.25 mM + NaCl 50 mM) and used to prepare lysis mix (1 ml lysis buffer + 100 µl 10% SDS + 10 µl proteinase K 18.6 mg/ml). Pellet with whole-blood leucocytes was thawed on ice and resuspended in 1 ml lysis mix. After 3h-incubation in the thermoblock at 55°C with agitation, NaCl was added to a final concentration of 1.5M. The tube was centrifuged at maximum speed for 10 min at RT, and supernatant was transferred to a new tube. Cold 100% isopropanol was added and pipetted up and down for proper homogenization. The tube was incubated overnight at -20°C and centrifuged at maximum speed for 15 min at 4°C. The supernatant was discarded, and the pellet was resuspended in 70% ethanol. The tube was centrifuged again at maximum speed for 15 min at 4°C, the supernatant was discarded, and the pellet was dried at RT until complete evaporation of ethanol. DNA was resuspended in Tris-EDTA (TE) buffer and incubated in the thermoblock for 10 min at 55°C with agitation for proper rehydration. Quality and quantity of DNA were evaluated using 2200 TapeStation (Agilent®) and Nanodrop One spectrophotometer (ThermoFisher Scientific®).

### DNA extraction from urine

Freshly-collected urine was transferred to 50 ml tubes and centrifuged at 3,000*g* for 10 min at RT. Supernatant was discarded, and pellet was washed in PBS prior to storage at -80°C. DNA extraction was performed as for whole-blood leucocytes.

### DNA extraction from saliva

Saliva was freshly-collected into Oragene 500 DNA tubes, and DNA was extracted according to manufacturer’s instructions (DNA Genotek). In brief, the tube was incubated in an air incubator overnight at 50°C, and the content was transferred to a 15 ml tube. 1/25 volumes of PT-L2P® (DNA Genotek) were added, and the tube was vortexed and incubated on ice for 10 min. The tube was centrifuged at maximum speed for 10 min at RT, and the supernatant was transferred to a new 15 ml tube. 1.2x volumes of 100% ethanol were added, and the sample was gently mixed and incubated for 10 min at RT to allow complete DNA precipitation. The tube was centrifuged at maximum speed for 10 min at RT, the supernatant was discarded, and the pellet was washed with 1 ml of 70% ethanol for 1 minute. DNA was rehydrated with 500 µl TE buffer and quantified using Qubit® BR kit on a Qubit®2.0 fluorimeter (ThermoFisher Scientific®), and DNA integrity was measured through 2200 Tapestation (Agilent®).

### Whole Genome Sequencing

Whole Genome Sequencing (WGS) was performed using DNA extracted from M116’s blood, saliva, and urine samples. DNA was fragmented to 350 bp, followed by library preparation, paired-end 150 bp WGS, and variant calling, all conducted by Novogene®. Sequencing was carried out with a 10X coverage (approximately 60 Gb and 200 million reads per sample) using the NovaSeq X Plus platform (Illumina). After quality control performed by Novogene (QD<2.0, FS>60.0, MQ<40.0, HaplotypeScore>13.0, MappingQualityRankSum<−12.5, and ReadPosRankSum<−8.0), approximately 3.8 million single nucleotide variants (SNVs) were identified. Variants were further filtered to include only those present in at least 2 out of the 3 samples, thereby minimizing the impact of technical artifacts. A control cohort consisting of 75 women from the Iberian population in Spain, was used for comparison. The same filters applied to the M116 samples were used for this cohort. All samples were processed and analyzed uniformly.

SNVs were annotated using the Ensembl Variant Effect Predictor (VEP) to determine transcript location, variant class, and additional attributes such as allele frequency (AF) from various populations, including 1000 Genomes Project (1000G) and gnomAD EUR exomes. Functional variant effect annotations were obtained from SIFT, PolyPhen-2, and Combined Annotation-Dependent Depletion (CADD) scores. Rare variants were defined as those with an AF < 0.015 in both the 1000G 30X Illumina NovaSeq sequencing data set (2,504 unrelated individuals) and gnomAD v4.1 genome data set. These rare variants were further classified based on their potential impact on protein function or their predicted effects according to sequence annotations, and were designated as variants of interest (VOIs). To compare the rarity of M116’s rare variants with the control cohort, three burden tests were performed: Cohort Allelic Sum Test (CAST), Rare Variant Test (RVT), and Rank Test of Proportions. Rare variants were considered differentially enriched if p-value < 0.05 and significant in at least 2 out of the 3 tests. Functional analyses, including over-representation analysis, were conducted using genes carrying VOIs or differentiating genes. These analyses employed ontology gene sets from the Human MSigDB collection. Pathways were deemed over-represented with a p-value < 0.05.

### Mitochondrial analyses

PBMCs from supercentenarian and healthy women controls (obtained from Banc de Sang i Teixits, Barcelona, Spain) were washed and plated in 96 well plates (25,000 cells/well) in RPMI (Gibco®, Cat. 11875093). PBMCs were simultaneously stained with 100 nM MitoTracker Green (Invitrogen®, Cat. M7514) for 30 min at RT together with either 50 nM Red Tetramethylrhodamine, ethyl ester (TMRE) (Invitrogen®, Cat. T669) for 30 min at RT, 5 µM MitoSOX Red (Invitrogen®, Cat. M36008) for 10 min at 37°C, or 7.5 µM BioTracker ATP-Red (Sigma-Aldrich®, Cat. SCT045) for 15 min at 37°C following manufacturer’s instructions. Cells were washed with PBS and analyzed by flow cytometry (FACSCanto II, BD Biosciences®) using FITC (MitoTracker Green) and PE (Red TMRE, MitoSOX Red, and BioTracker ATP-Red) channels. Only MitoTracker Green positive cells were considered in the subsequent analytical assessments. Unpaired t-test was performed to statistically compare M116 to the mean of controls.

### Clonal Hematopoiesis Analysis

Genomic DNA was obtained from peripheral blood samples and was used for targeted deep sequencing (TDS) studies. Barcoded libraries were prepared according to the manufacturer’s instructions, using a probe-based panel (KAPA HyperCap, Roche®) targeting frequently mutated regions of 50 myeloid-related genes (**Data set 6**). Samples were run on a MiSeq (Illumina®) sequencer for paired-end 2×75 bp reads with a mean coverage of 1000X. Sequencing data were analyzed using an in-house pipeline. Reads were aligned to human genome build 19 (hg19/GRCh37) using BWA 0.7.15. Post-alignment and base recalibration were performed using the tools in GATK 4.1.8.0 software package. Variant calling was performed with Mutect2 4.1.8.0, which is included in GATK software package. ANNOVAR 20200607 was used for variant annotation. The variant filtering process was based on the criteria proposed by the Spanish Group of myelodysplastic syndromes. Briefly, variants were filtered according to location (exonic and splicing), variant type (nonsynonymous single-nucleotide variants and small insertions/deletions), read depth (>100x), minor allele frequency (MAF < 0.01 according to dbSNP, ExAC, Exome Variant Server and 1000 Genomes project population databases) and variant allele frequency (VAF ≥ 2%).

### Single-cell transcriptomics

scRNA-seq was performed using Chromium Next GEM Single Cell 3’ Kit v3.1 (10X Genomics) according to the manufacturer’s instructions. Briefly, PBMCs were thawed and quantified in order to calculate the number of cells to be loaded. Then, barcoded Single Cell 3ʹ Gel Beads, a master mix containing PBMCs, and Partitioning Oil were combined onto the Chromium Next GEM Chip G to generate Gel Beads-In-Emulsion (GEMs), where polyadenylated mRNAs were reverse transcribed. This way, all generated cDNAs from the same cell shared a common 10X Barcode. Following the reverse transcription, GEMs were broken and cDNAs were amplified and cleaned up with SPRIselect beads (Beckman Coulter®). Next, a portion of these cDNAs were enzymatically fragmented and subjected to adaptor ligation before using them as a PCR template for the incorporation of i5/i7 indexes. Finally, libraries were purified with SPRIselect beads, quantified and quality checked by using the 2200 TapeStation (Agilent®), and subjected to paired-end 150-bp sequencing (Novaseq systems, Illumina®) following standard practices at external facilities (Novogene®).

The gene count matrices were analyzed using the Seurat package (version 5.1.0) in R (version 4.4.1). Quality control (QC) was performed to ensure that high-quality cells were retained for downstream analysis. Genes expressed in less than 3 cells were filtered out. Mitochondrial gene expression was quantified using PercentageFeatureSet function, and cells with mitochondrial content >20% were excluded. Cells with less than 300 detected genes were also filtered out as low quality. To exclude potential artificial doublets generated during library construction, we identified them applying two independent methods: DoubletFinder (version 2.0.4) and scDblFinder (version 1.20.0). Cells classified as doublets by both methods were filtered out for downstream analysis. Data was normalized (NormalizeData function: “LogNormalize” method), followed by dimensionality reduction performed via principal component analysis (RunPCA function) on the scaled expression (ScaleData function) of the top 2,000 highly variable genes (FindVariableFeatures function: variance-stabilizing transformation (VST) method). 15 principal components (PCs) were retained for downstream analysis, by inspecting the elbow plots. These top PCs were used to create a UMAP embedding (RunUMAP function) and cluster the cells (FindClusters function) in a 20-nearest neighbor graph (FindNeighbors function). Cell type annotations were predicted with automatic label transfer using CellTypist (version 1.6.3) (with the human Immune_All_High or Immune_All_Low models). This study analyzed two publicly available scRNA-seq data sets from human PBMCs: Terekhova et al., Synapse database under accession code syn49637038; and at http://gerg.gsc.riken.jp/SC2018/. Normalized data was subsampled to retain 10,000 cells from each age group. In both data sets, CellTypist was used to annotate the cells, as previously described.

### Metabolome

300 µl of serum sample was used for the whole proton nuclear magnetic resonance (^1^H-NMR) analysis, which includes the lipoprotein profile based on the Liposcale test, the glycoprotein profile and the Low Molecular Weight Metabolites (LMWM) profile from intact serum ^1^H-NMR spectra; and the specific lipid species characterization, from the lipid serum extract ^1^H-NMR spectra. High-resolution ^1^H-NMR spectroscopy data was acquired on a Bruker 600 MHz Spectrometer: 1D Nuclear Overhauser Effect Spectroscopy (NOESY), Carr-Purcell-Meiboom-Gill (CPMG) was used to characterize small molecules such as aminoacids and sugars; and LED Diffusion (Diff) experiments, to detect larger molecules such as lipoproteins, glycoproteins and choline compounds. All the sequences were run at 37°C in quantitative conditions. We obtained the lipid extract using a biphasic extraction with methanol/methyl-tert-butyl ether. For NMR measurements, the lipid extract was dried and reconstituted in 0.01% tetramethylsilane (TMS) solution (0.067 mM) and deuterated solvents. From intact serum, the Liposcale test (IVD-CE marked) was used to obtain the composition, the mean size and the number of lipoproteins particles of nine subtypes (large, medium and small) of the main lipoprotein types (VLDL, LDL and HDL). From the same NMR spectra, we obtained the general measurement of circulating glycoproteins by deconvoluting with analytical functions the specific region where glycoproteins resonate, to quantify the area, proportional to the concentration of the acetyl groups of N-acetylglucosamine and N-acetylgalactosamine (GlycA) and acetyl groups of N-acetylneuraminic acid (GlycB). Complementary, we applied a CPMG filter to profile the ^1^H-NMR spectra to obtain the concentration of a set of LMWM such as aminoacids and sugars from the same serum sample. Finally, we obtained the concentration of the major lipid classes (fatty acids, glycerolipids, phospholipids and sterols) and some individual species from the ^1^H-NMR spectra of the previously extracted serum sample using the BUME protocol. Lipid quantification, based on the Lipspin software, relied on lineshape fitting analysis of spectral regions. Since the spectral area is equivalent to the molecular abundance, individual signal areas were converted into molar concentrations by normalizing by external measurements. The lipid species obtained by this NMR approach included: cholesterol (free and esterified), unsaturated fatty acids (omega-6, omega-7, omega-9, omega-3), saturated fatty acids, monounsaturated fatty acids, linoleic acid, docosahexaenoic acid, arachidonic acid and eicosapentaenoic acid; glycerides and phospholipids (total cholines, triglycerides, phosphoglycerides, phosphatidylcholine, sphingomyeline and lysophosphatidylcholine).

The concentration of metabolites was compared with the population values (median, 25-75 percentiles) using the same methodological approach and NMR equipment. Specifically, the lipoprotein and glycoprotein profiles were compared with the general population values from a total of 6,022 individuals across two Spanish cohorts: the Study and the Mollerussa Study. These are population-based Spanish cohorts comprising individuals aged 18 years and older, with a composition of 55% women. Similarly, the concentration of the aqueous metabolome and lipid families were compared with the general population values from a total of a subset of 1,965 individuals from the study (57% women).

### Proteomics

100 µl plasma of each sample was pre-cleared by centrifugation at 3000*g* for 10 min. For extracellular vesicles (ECV) extraction, 1M ammonium acetate was added to precipitate ECVs on ice for 45 min. Then, 100 mM ammonium acetate was added to the mixture, and ECVs were precipitated by centrifugation at 20,000*g* for 30 min. ECVs were washed with 50 mM ammonium bicarbonate (Sigma-Aldrich). Then, 600 µl of 1% ammonium deoxycholate (Sigma Aldrich) were added. The concentration of protein in each sample was measured using a bicinchoninic acid assay (BCA assay). Ammonium bicarbonate was used to dilute 500 µg of protein into a final volume of 500 µl. Next, dithiothreitol (Sigma-Aldrich) was added to obtain a final concentration of 20 mM, followed by iodoacetamide (Sigma-Aldrich) to a final concentration of 40 mM. Next, trypsin (Roche) was added to the sample in a 1:25 protein ratio and incubated at 37°C overnight. Next day, formic acid (Fisher scientific) in a final concentration of 0.1% was added and extraction of proteins was done using Empore™ Solid Phase Extraction Cartridges (3M), following manufacturer’s instructions. The eluted samples were then centrifuged for 90 min using a speed vacuum centrifuge (Thermo, RC1010), followed by snap freezing in liquid nitrogen. Then, the samples were kept in a freeze dryer (LyoDry Compact Benchtop, MechaTech) overnight. Next, the samples were reconstituted in 30 µl of 0.1% formic acid (FA) and an o-Phthaladehyde (Oparil) assay was performed to determine the concentration of each sample. After that, the sample was prepared in a concentration of 0.5 µg/µl using 0.1% FA and alcohol dehydrogenase (ADH). The samples were prepared in glass mass spectrometry vials for proteomic analysis using a Waters Synapt G2Si High-Definition Mass Spectrometry (Waters Corporation) operated by the MassLynx 4.1., 110 min running time with 2 µl of an injection containing 1 µg of peptide. Quality controls were also run along with samples to guarantee consistency. Pooled quality controls were made from all samples, in which the samples were run at the beginning, middle and end of the mass spectrometry run. Samples were randomized before running the experiment. The proteomic data was then imported into Progenesis software 4.2 (Nonlinear Dynamic, UK) to identify and quantify peptides and proteins.

Data were analysed in R Studio. The intensity data protein abundances were normalized using Trimmed Mean of M-values (TMM) normalization using the NOISeq package. Technical replicates effects were corrected by using ARSyNseq, and log2 transformation was applied to stabilize variance. Orthogonal Partial Least Squares Discriminant Analysis (OPLS-DA) was performed using ropls package to identify group separations. For differential expression analysis, the limma package was used. Statistical significance was assessed using empirical Bayes moderation (eBayes), with differentially expressed proteins identified based on an adjusted p-value < 0.05 and |logFC| > 1.

To identify Biological Processes (BP) associated with differentially expressed proteins, Gene Ontology (GO) enrichment analysis was performed using clusterProfiler. The enrichGO function was applied selecting the BPs GO terms with a p-value and q-value cutoff of 0.05. To classify GO terms into general BP clusters, Wang’s semantic similarity method was used with the GOSemSim package to calculate term similarities. A heatmap was created to visualize clustering, which was further redefined into eight functional clusters. A Circos plot was used to visualize the relationships between differentially expressed genes and the GO clusters, using the circlize package.

### 16S meta-genomics sequencing

Stool was collected from 3 different days (3 biological replicates) and kept at -20°C until DNA extraction (Biobanc IDIBGI and Goodgut®). Total genomic DNA was extracted from 150–200 mg of each stool sample after homogenisation using the DNeasy Powersoil Pro kit (Qiagen®) according to manufacturer’s instructions. Quality and quantity of DNA were evaluated using Qubit® BR kit on a Qubit®2.0 fluorimeter (ThermoFisher Scientific®) and on a Nanodrop ND-2000 UV-Vis spectrophotometer (ThermoFisher Scientific®). The v3-v4 region of the bacterial 16S rRNA gene was amplified and sequenced (paired-end 250-bp) following standard practices at external facilities (Novogene®) using primers 515F (5’-GTG CCA GCM GCC GCG GTA A-3’) and 806R (5’-GGA CTA CHV GGG TWT CTA AT-3’).

Obtained reads were processed with the dada2 pipeline and phyloseq R package. Default settings were used for filtering and trimming reads. Built-in training models were used to learn error rates for the amplicon data set. Identical sequencing reads were combined through dada2’s dereplication functionality, and the dada2 sequence-variant inference algorithm was applied to each data set. Subsequently, paired-end reads were merged, a sequence table was constructed, taxonomy was assigned, and abundance was calculated at all possible taxonomic levels using the Silva’s 138 dada2-formatted database (www.arb-silva.de). Diversity indices (i.e., Chao1 and Shannon) together with beta-diversity matrices (e.g., weighted unifrac and principal coordinate analyses (PCoA)) were computed using phyloseq, vegan, ape, MicrobiotaProcess, UpsetR and mia R packages.

As independent validation, we considered the publicly available data sets collected and curated in the curatedMetagenomicData R package. Of the available data sets, we selected those comprising samples matching the following criteria: (i) gut samples collected from healthy individuals (“disease == healthy”); (ii) samples with available age value and classified as senior (“age_category == senior”), which comprised between 65 and 91 years old; and finally (iii) those samples without antibiotic use on the day of sampling (available information; antibiotics_current_use !=“yes”). A total of 445 samples were analyzed from the curatedMetagenomicData data set.

### Infinium MethylationEPIC BeadChip

Genome-wide DNA methylation profiling was performed at the Genomics Unit of the Josep Carreras Leukaemia Research Institute. Briefly, DNA samples were quantified with Qubit® BR kit on a Qubit®2.0 fluorimeter (ThermoFisher Scientific®), and integrity was checked using agarose gel electrophoresis. DNA from samples passing the quality control was bisulfite-converted using the EZ DNA Methylation-Gold™ Kit (Zymo Research) following the manufacturer’s instructions. Then, bisulfite-converted DNA was hybridized into the Methylation EPICv1.0 BeadChip (Illumina) array interrogating > 850,000 CpG sites according to manufacturer’s instructions as previously described^37^. Fluorescent signal was detected by the Illumina iScan confocal laser scanner with the Autoloader system. Illumina methylation raw data (idat files) were preprocessed using the R environment (version 4.3.2) with the minfi package (version 1.48.0). Raw signal intensities were normalized applying a background correction method with dye-bias normalization using ssNoob method. Then, probes with a detection p-value > 0.01, cross-reacting probes and probes that overlapped single nucleotide variants and sex-chromosomal probes were removed as quality control steps. The DNA methylation levels were represented as β-values ranging from 0 to 1 (corresponding to 0-100% methylation). Supervised methylation analysis was performed comparing the difference between M116 saliva samples to the mean methylation of the control cohort hybridized and pre-processed as mentioned above^38^. Differentially methylated CpGs were considered if Δβ > |0.50| as stated in the manuscript. Differentially methylated CpG probes were used to perform hierarchical clustering analysis with canberra distance and ward.D2 clustering method. The same CpG probes were used for the t-SNE analysis. To correlate methylation and expression values, the common CpGs differentially methylated located within a gene were selected in a panel of human cell lines and correlated with the expression of the same gene^40^. We considered a significant correlation between methylation and expression if p-value < 0.05 and Spearman Rho > |0.3|. To determine the methylation status in repetitive elements, the REMP R package was used. The methylation values in repetitive elements were predicted for ALU, LINE-1 and ERV applying the default parameters, using a random forest model for the prediction with 1,000 bp windows within the genome. The median methylation level of each repetitive element for individual was represented in a boxplot. Finally, epigenetic clocks, with the exception of the ribosomal DNA methylation clock, were calculated using the methylclock package in R following the recommended pipeline.

### Whole Genome Bisulfite Sequencing

DNA fragmentation (350 bp), library preparation, bisulfite conversion, and paired-end-150 whole genome bisulfite sequencing was conducted by Novogene®. Sequencing was performed as for Whole Genome Sequencing. rDNA methylation age estimates were calculated as previously described (Wang and Lemos, 2019) and compared to individuals from the Reprocell biobank repository, with no known health condition.

## Supporting information

Data Set S1

Data Set S2

Data Set S3

Data Set S4

Data Set S5

Data Set S6

Data Set S7

Data Set S8

Data Set S9

Data Set S10

Data Set S11

Data Set S12

## Data availability

All data supporting the findings of this study are available within the source data provided with this article. Multiomics data have been deposited in the European Genome-Phenome Archive (EGA), under restricted acces to protect individual information. Access will be granted for appropiate use for researchers by the corresponding author.

## Code availability

There was no new code created for the analysis of these data.

## Acknowledgements

Research in ME group is funded by the CERCA Programme / Generalitat de Catalunya; MCIN/ AEI /10.13039/501100011033/ and the European Regional Development Fund, ‘A way to make Europe’ ERDF (project PID2021-125282OB-I00); Departament de Recerca i Universitats / Generalitat de Catalunya (2021 SGR 01494), European Union under THRIVE Grant Agreement Nr. 101136622, “La Caixa” Research Foundation and the Cellex Foundation (CEL007). ESP is a fellow of the Spanish Ministry of Science, Innovation and Universities, under FPI contract no. PRE2022-105015. GF is recipient of Ayuda Investigador AECC 2023 (INVES234765FERR), Fundación Científica AECC. CQD is a fellow of the Spanish Ministry of Science, Innovation and Universities, under FPU contract no. FPU22/01655. Research in AN group is supported by the I+D+i project PID2021-127792NB-I00 funded by MCIN/AEI/10.13039/501100011033 (FEDER Una manera de hacer Europa); and “Unidad de Excelencia María de Maeztu,” funded by the AEI [CEX2018-000792-M] and Departament de Recerca i Universitats de la Generalitat de Catalunya [GRC 2021 SGR 0467]. Research in MDM group is funded by the Departament de Recerca i Universitats / Generalitat de Catalunya (2021 SGR 01366). Research in SM group was supported by the was supported by the Instituto de Salud Carlos III (PI20/00328) and the M. C. Andreu Memorial Fund. JR-B was supported by a doctoral grant from the Universitat Oberta de Catalunya. Research in MB lab is funded by the European Union, project ERC-AvG Shelterins (GA 882385), Horizon 2020 Programme. The mass spectrometry work was supported by the John and Lucille van Geest Foundation, and the National Institute for Health and Care Research Leicester Biomedical Research Centre. ME is an ICREA Research Professor. Special thanks are given to personnel taking care of M116 and to all the biobanks’ technicians involved and contributing to keep samples and procedures for the current research with special mention to Gerard Pardo Albiñana.

## Author contributions

ESP, ANC, MCP, CAGP, ICM, CQD, ECG, ABC, FS, GF, and VD: conceptualization, performing experiments, formal analysis and writing. EM: scRNA-seq analysis supervision. RP and CA: DNA methylation experiments. LS: genetic analysis. CDLT: proteomics analysis. FV: statistics. IG: cytogenetic analysis. NC: sample collection. PA, FS and MM: genetic analysis. CM: scRNA-seq analysis supervision. SP, TG, ML, MP, RCT, AL, LJGG, and XA: microbiome analysis. SS, PT, JR, MG, NA, and OY: metabolome analysis. PM, RSV, and MAB: telomeric analysis. JO and BL: rDNAm analysis. JRB, MT, MMS, KAK, THC, PAQ, DJLJ, and SM: proteomics analysis. CV, EBB, MM, FB, GM, HL, and AN: genetic analysis. ME: conceptualization, writing and scientific supervision.

