## Supplementary material for "The Multiomics Blueprint of Extreme Human Lifespan": Figure S1

### Slide 1
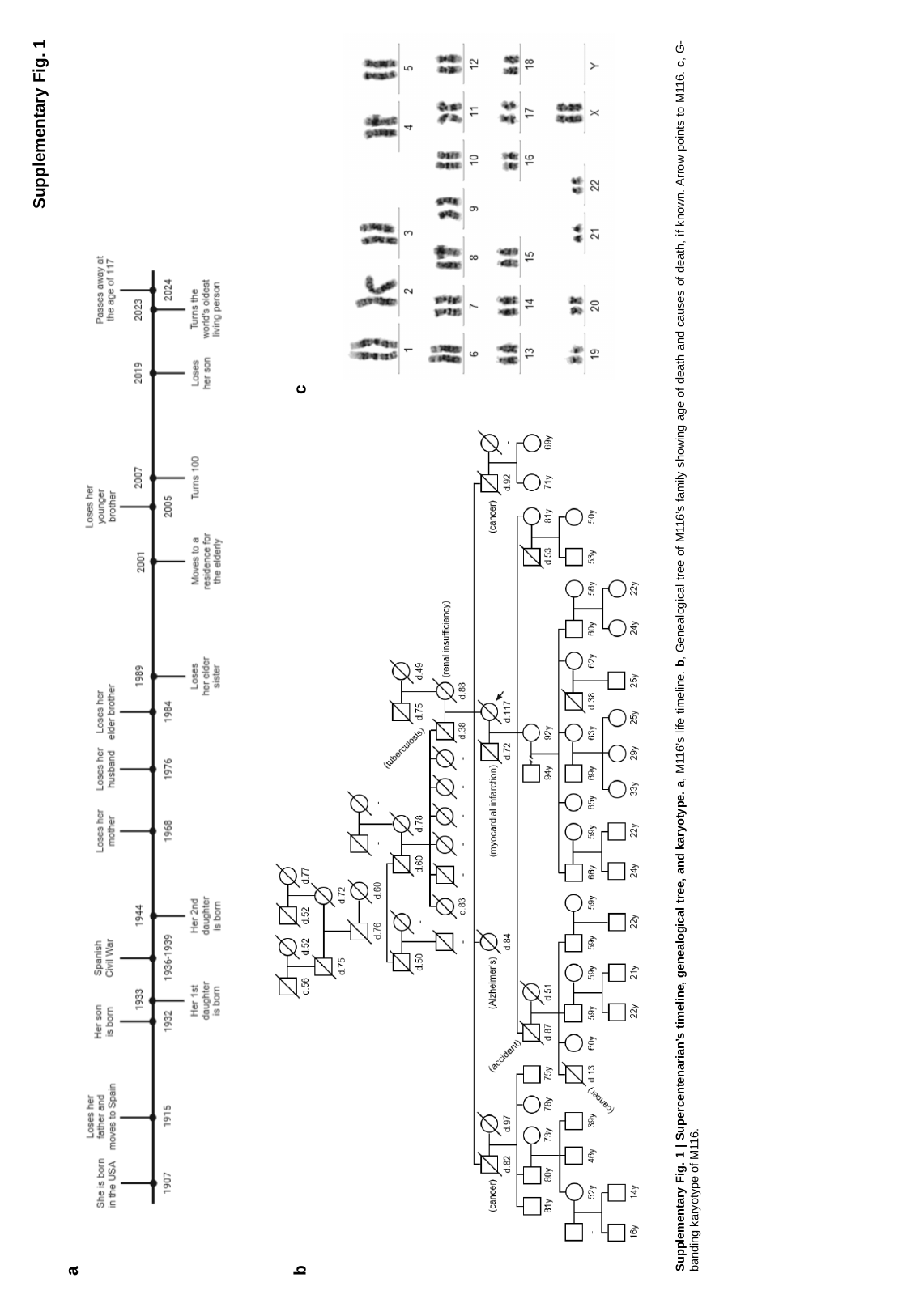

b
a
Supplementary Fig. 1 | Supercentenarian’s timeline, genealogical tree, and karyotype. a, M116’s life timeline. b, Genealogical tree of M116’s family showing age of death and causes of death, if known. Arrow points to M116. c, G-banding karyotype of M116.
c
Supplementary Fig. 1
