## Supplementary material for "The Multiomics Blueprint of Extreme Human Lifespan": Figure S2

### Slide 1
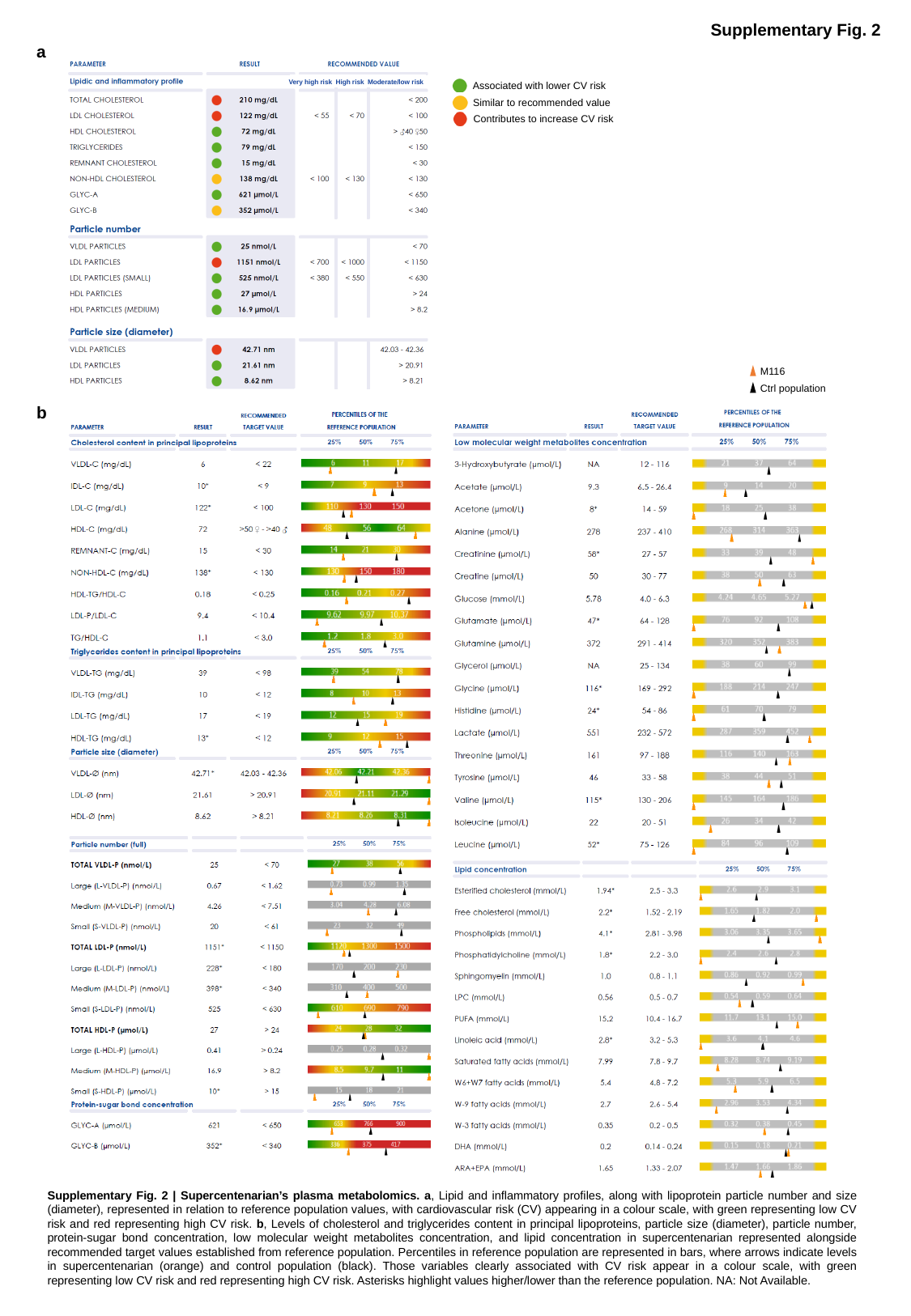

Supplementary Fig. 2
a
Very high risk High risk Moderate/low risk
Associated with lower CV risk
Similar to recommended value
Contributes to increase CV risk
M116
Ctrl population
b
Supplementary Fig. 2 | Supercentenarian’s plasma metabolomics. a, Lipid and inflammatory profiles, along with lipoprotein particle number and size (diameter), represented in relation to reference population values, with cardiovascular risk (CV) appearing in a colour scale, with green representing low CV risk and red representing high CV risk. b, Levels of cholesterol and triglycerides content in principal lipoproteins, particle size (diameter), particle number, protein-sugar bond concentration, low molecular weight metabolites concentration, and lipid concentration in supercentenarian represented alongside recommended target values established from reference population. Percentiles in reference population are represented in bars, where arrows indicate levels in supercentenarian (orange) and control population (black). Those variables clearly associated with CV risk appear in a colour scale, with green representing low CV risk and red representing high CV risk. Asterisks highlight values higher/lower than the reference population. NA: Not Available.
