## Supplementary material for "The Multiomics Blueprint of Extreme Human Lifespan": Figure S3

### Slide 1
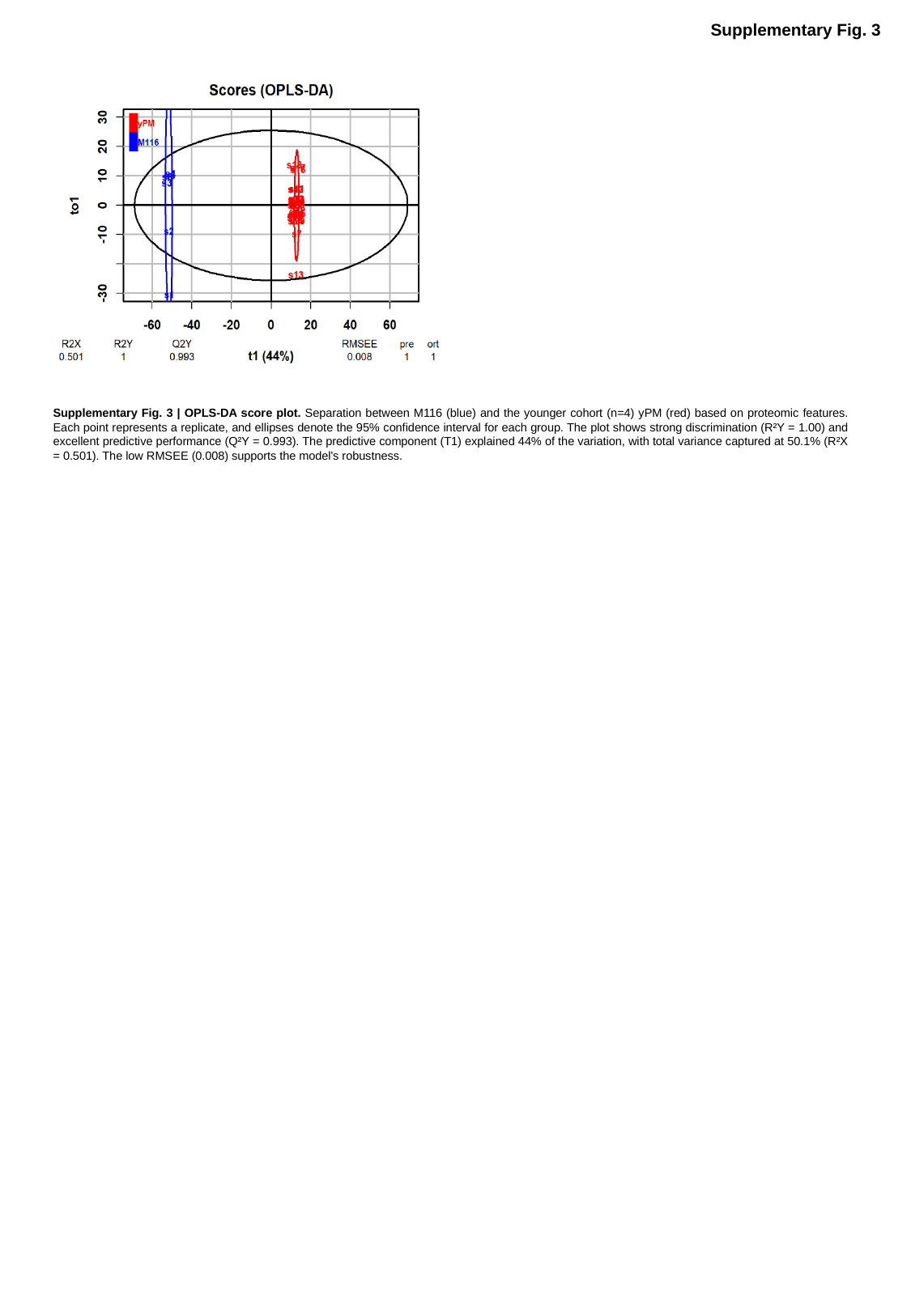

Supplementary Fig. 3
Supplementary Fig. 3 | OPLS-DA score plot. Separation between M116 (blue) and the younger cohort (n=4) yPM (red) based on proteomic features. Each point represents a replicate, and ellipses denote the 95% confidence interval for each group. The plot shows strong discrimination (R²Y = 1.00) and excellent predictive performance (Q²Y = 0.993). The predictive component (T1) explained 44% of the variation, with total variance captured at 50.1% (R²X = 0.501). The low RMSEE (0.008) supports the model's robustness.
