## Supplementary material for "The Multiomics Blueprint of Extreme Human Lifespan": Figure S4

### Slide 1
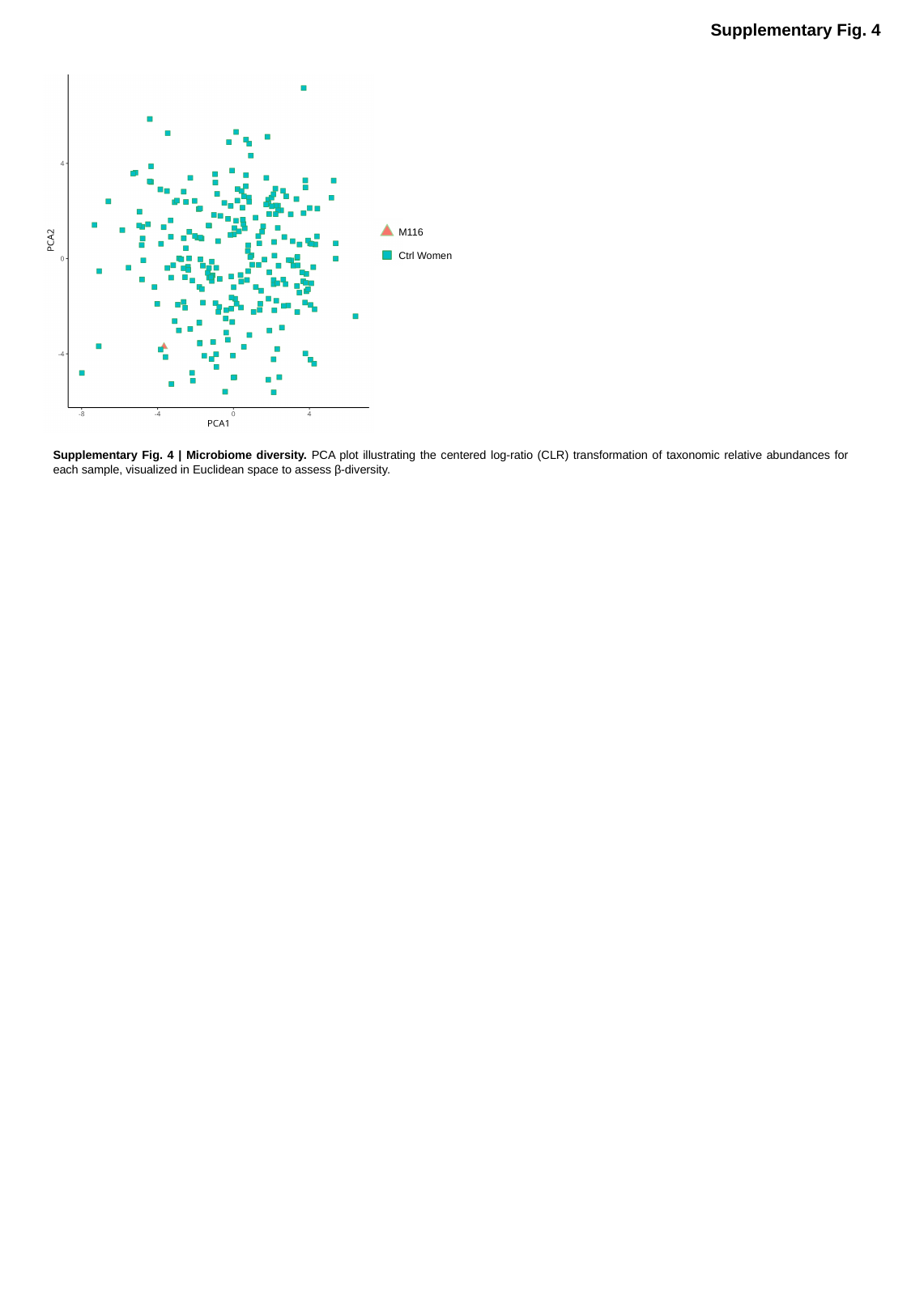

Supplementary Fig. 4
M116
Ctrl Women
Supplementary Fig. 4 | Microbiome diversity. PCA plot illustrating the centered log-ratio (CLR) transformation of taxonomic relative abundances for each sample, visualized in Euclidean space to assess β-diversity.
