## Supplementary material for "The Multiomics Blueprint of Extreme Human Lifespan": Figure S5

### Slide 1
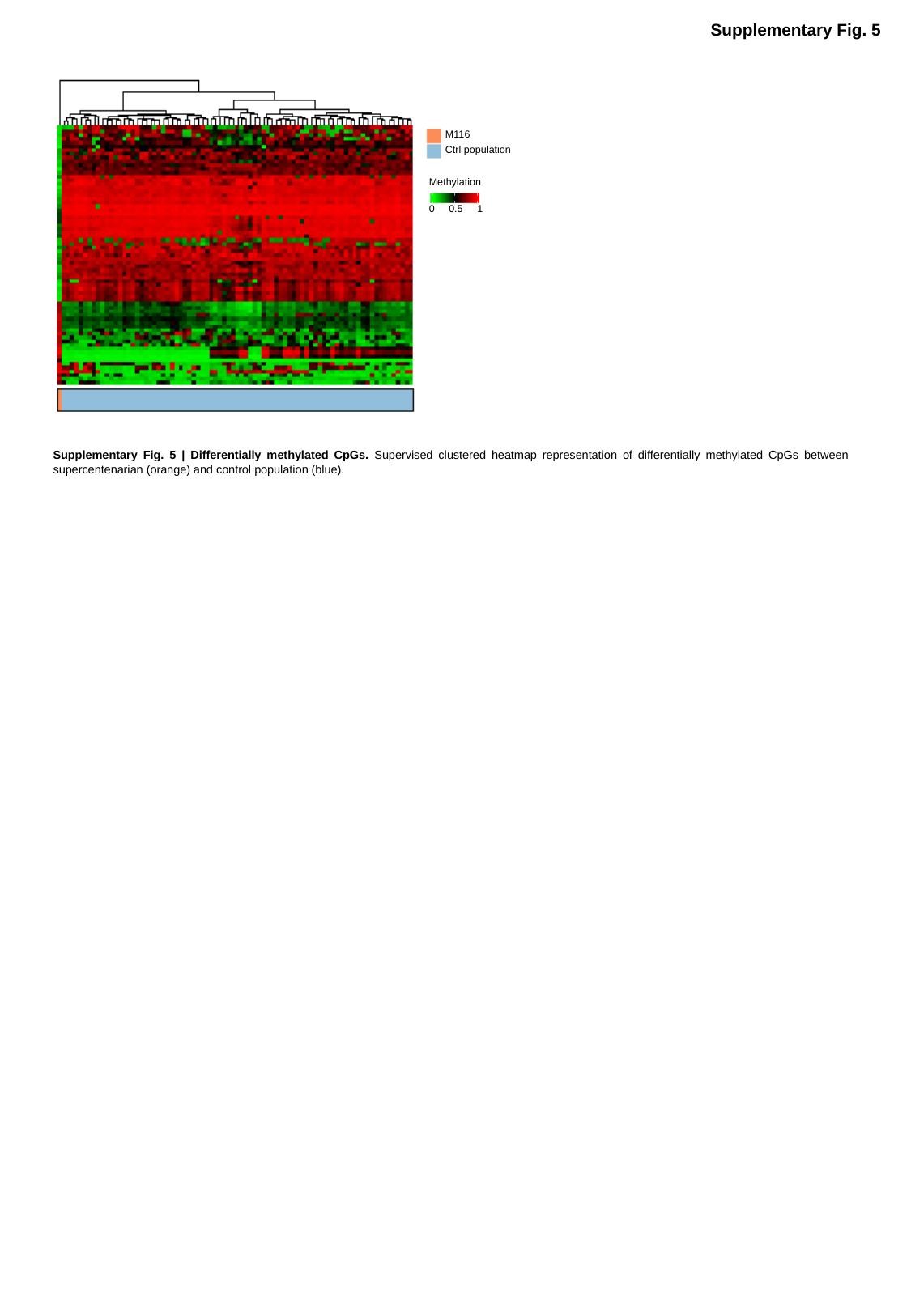

Supplementary Fig. 5
M116
Ctrl population
Methylation
0 0.5 1
Supplementary Fig. 5 | Differentially methylated CpGs. Supervised clustered heatmap representation of differentially methylated CpGs between supercentenarian (orange) and control population (blue).
